# Genetic differentiation is determined by geographic distance in *Clarkia pulchella*

**DOI:** 10.1101/374454

**Authors:** Megan Bontrager, Amy L. Angert

**Affiliations:** Department of Botany, University of British Columbia, Vancouver, British Columbia V6T 1Z4, Canada; Departments of Botany and Zoology, University of British Columbia, Vancouver, British Columbia V6T 1Z4, Canada

**Keywords:** *Clarkia*, isolation-by-distance, landscape genetics, population genetics, genetic diversity

## Abstract

Both environmental differences and geographic distances may contribute to the genetic differentiation of populations on the landscape. Understanding the relative importance of these drivers is of particular interest in the context of geographic range limits, as both swamping gene flow and lack of genetic diversity are hypothesized causes of range limits. We investigated the landscape genetic structure of 32 populations of the annual wildflower *Clarkia pulchella* from across the species’ geographic range in the interior Pacific North-west. We tested whether climatic differences between populations influenced the magnitude of their genetic differentiation. We also investigated patterns of population structure and geographic gradients in genetic diversity. Contrary to our expectations, we found an increase in genetic diversity near the species’ northern range edge. We found no notable contribution of climatic differences to genetic differentiation, indicating that any processes that might operate to differentiate populations based on temperature or precipitation are not affecting the putatively neutral loci in these analyses. Rather, these results support seed and pollen movement at limited distances relative to the species’ range and that this movement and the subsequent incorporation of immigrants into the local gene pool are not influenced by temperature or precipitation similarities among populations. We found that populations in the northern and southern parts of the range tended to belong to distinct genetic groups and that central and eastern populations were admixed between these two groups. This pattern could be the result of a past or current geographic barrier associated with the Columbia Plateau, or it could be the result of spread from separate sets of refugia after the last glacial maximum.

## Introduction

Geographic distance is often a primary predictor of genetic differentiation among populations on the landscape. Populations that are near each other are often more genetically similar, while distant populations are often more divergent. This pattern arises when the dispersal distances of individuals and gametes are small relative to the distances separating populations; as a result, differences accumulate among populations due to drift faster than they are homogenized by gene flow (Slatkin, 1993; Wright, 1943). Isolation by distance is well-documented and prevalent (Sexton et al., 2014) to the extent that it is a reasonable null expectation for how genetic differentiation is structured at geographic scales.

However, geographic distance is not the only factor that structures dispersal and realized gene flow among populations (McRae, 2006; Epps et al., 2005). Not all geographic distances are equivalent in the extent to which they might facilitate or impede gene flow (Storfer et al., 2007). Landscape features between populations may impose barriers to gene flow beyond those predicted by geographic distance. Gaps in suitable habitat may be large enough that very few instances of gene flow occur across them, leading to differentiation of the populations on either side. For example, Reeves and Richards (2014) found genetic differentiation between populations of *Helianthus pumilus* that could be attributed to an unsuitable mountainous area interrupting the species’ distribution. Other features of the landscape might act as corridors for the organisms themselves or for agents of gene flow (i.e. seed dispersers or pollinators). For example, wind and water flow along rivers may increase gene flow among populations situated along them (Lee et al., 2018). In these types of scenarios we expect to see deviations from a strict pattern of isolation by distance, and population genetic structure will be better described by membership in discrete groups on either side of a barrier in the former case, or by patterns of admixture or increased similarity in populations connected by corridors in the latter.

Environmental differences between occupied sites may also contribute to the magnitude of genetic differentiation between populations (Slatkin, 1973; Wang and Bradburd, 2014). If populations are strongly locally adapted, then migrants that have moved between environments may be unable to survive to reproduction or may have low reproductive success (Nosil et al., 2005). In this case, realized gene flow may be low between different environments (Mosca et al., 2012). Similarly, vectors of gene flow such as pollinators and seed dispersers (or the organisms themselves, in the case of motile species) may have environmental preferences that lead to greater rates of gene flow among similar environments (Bolnick et al., 2009).

The current genetic structure of populations is also strongly influenced by past processes (Hewitt, 2004). In temperate regions including the Pacific Northwest, higher latitudes were glaciated until approximately 20,000 years ago (Booth et al., 2003) and this affected the distribution of many species, leaving lasting signatures on their genetic structure (Brunsfeld et al., 2001; Shafer et al., 2010). Species that previously had disjunct distributions—for example, those that occupied multiple refugia during glaciation—may exhibit multiple corresponding genetic clusters in the present day (Beatty and Provan, 2011; Carstens et al., 2013; Sproul et al., 2015). Populations that are the result of range expansions into previously glaciated areas may have lower levels of genetic diversity as a result of repeated founder events (Kuchta and Tan, 2005; Hewitt, 2004). These patterns may underlie (and sometimes confound) genetic structure that could also be attributed to isolation by distance or environment.

Despite the accumulation of numerous case studies, it is still challenging to draw generalizations about the extent to which the genetic structure of a given species is likely to be determined by geographic vs. environmental differences. A recent meta-analysis (Sexton et al., 2014) examined how the frequency of isolation by distance vs. by environment varied across broad taxonomic groups, and found that plants more frequently showed patterns of isolation by distance than vertebrates or invertebrates. However, in more than half of the plant species that displayed a pattern of isolation by distance, environmental similarity also contributed to genetic structure. In a small number of plant species, only environmental differences explained genetic structure. Although geography and environment may both have important effects on patterns of genetic differentiation, generalizations about when one will prevail over the other and what organismal traits determine their relative effect sizes remain elusive. The accumulation of more case studies and the development and use of more appropriate statistical methods will likely move this field forward (Wang and Bradburd, 2014; Bradburd et al., 2013).

The way that the landscape shapes genetic structure is of particular interest in the context of geographic range limits. Local adaptation may be constrained in range edge populations if these populations are inundated with gene flow from populations in dissimilar environments (Kirkpatrick and Barton, 1997). If populations are isolated by environmental differences, that might prevent swamping gene flow. Rather, gene flow between populations in similar environments could facilitate local adaptation by increasing adaptive genetic diversity (Sexton et al., 2011). This might be of particular importance if species occupy spatially heterogeneous environments, where random dispersal would otherwise result in frequent gene flow between divergent environments.

In this study, we investigated whether environmental differences between populations of the annual wildflower *Clarkia pulchella* contribute to their genetic differentiation, which we expected to also be strongly structured by geographic distances. Further, we explored whether patterns of genetic differentiation are better described by admixture among distinct genetic groups or continuous genetic differentiation across the landscape. We expected that topographic features, such as the Rocky Mountains, might be an impediment to the movement of seed dispersers and pollinators, and that this might result in disjunct genetic groups. Finally, we explored whether genetic diversity varies geographically in this species. We predicted lower levels of genetic diversity at high latitudes if this species has undergone a range expansion northward after the last glacial maximum.

## Methods

### Study species

*Clarkia pulchella* Pursh (Onagraceae) is a winter annual wildflower that grows east of the Cascade Mountains in the Pacific Northwest. It can be found in eastern Washington, eastern Oregon, Idaho, and western Montana (United States) and in southeastern British Columbia (Canada; Figure 1). It grows in large populations (i.e., thousands of flowering individuals) on open, south-facing slopes from 100 to 2200 meters elevation, though the majority of populations are found between 500 and 1600 m. While temperature generally decreases and precipitation generally increases from south to north and west to east across the range of *C. pulchella*, temperature and precipitation are also strongly influenced by elevation. Topographic complexity across the range creates large amounts of variation around geographic trends and appears to disrupt spatial autocorrelation in climate among populations of *C. pulchella* (Figure 2). This species has small seeds (c. 1 mm long) that lack an obvious dispersal mechanism. Flowers are visited by a diverse array of pollinators, including solitary bees, bee flies, bumblebees, and occasionally hummingbirds (M. Bontrager, personal observation).

**Figure 1.**
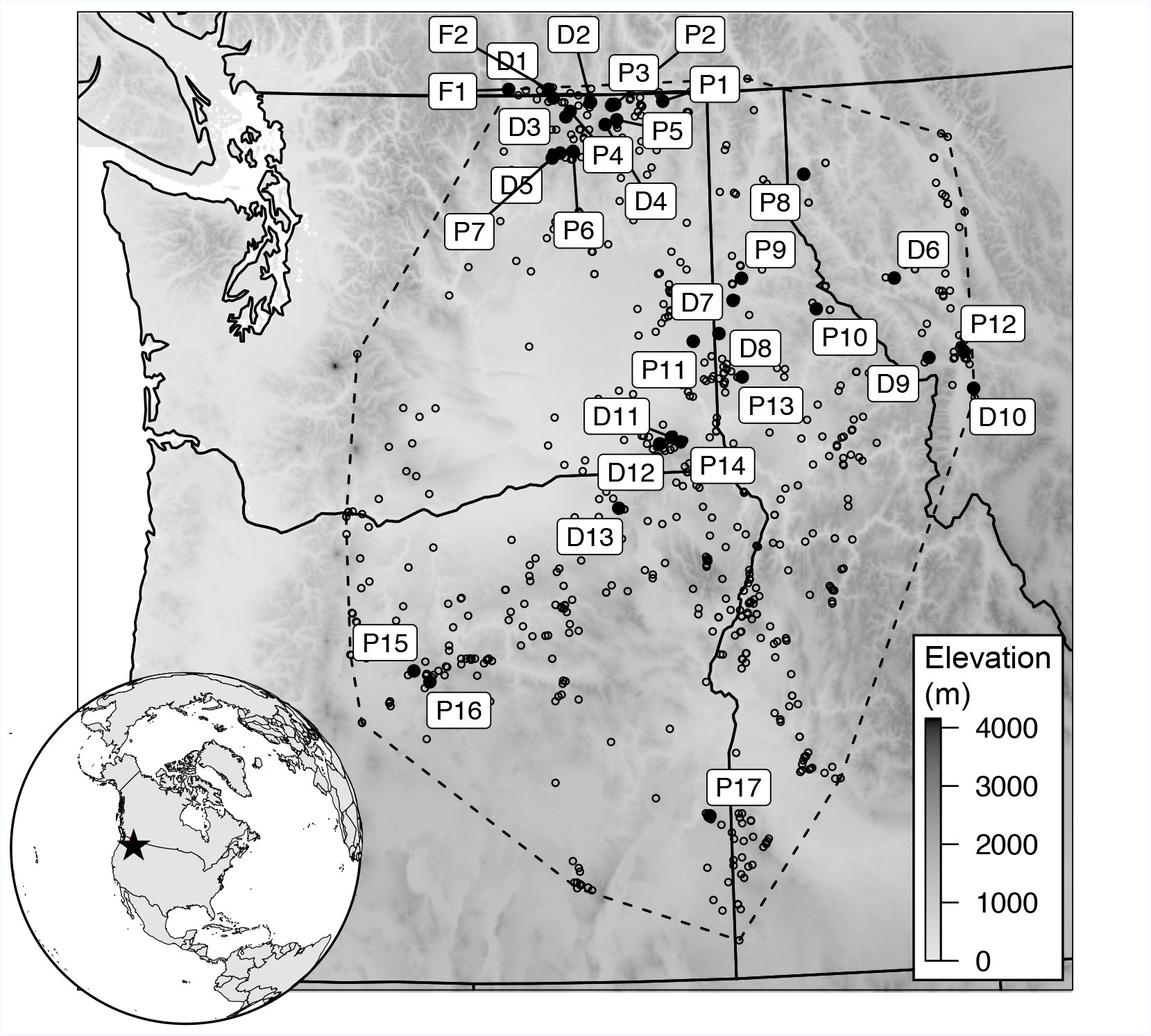
The geographic range of *Clarkia pulchella* across the interior of the Pacific Northwest. Small open points mark the locations of all herbarium records of *C. pulchella* from the Consortium of Pacific Northwest Herbaria that could be accurately assigned coordinates. The dashed line marks the maximum convex polygon drawn around these points. Larger filled points are populations that were sampled for this project. Labels correspond to population IDs in Table S1. Background shading shows elevation. The Columbia Basin is the unsampled area west of population D11.

**Figure 2.**
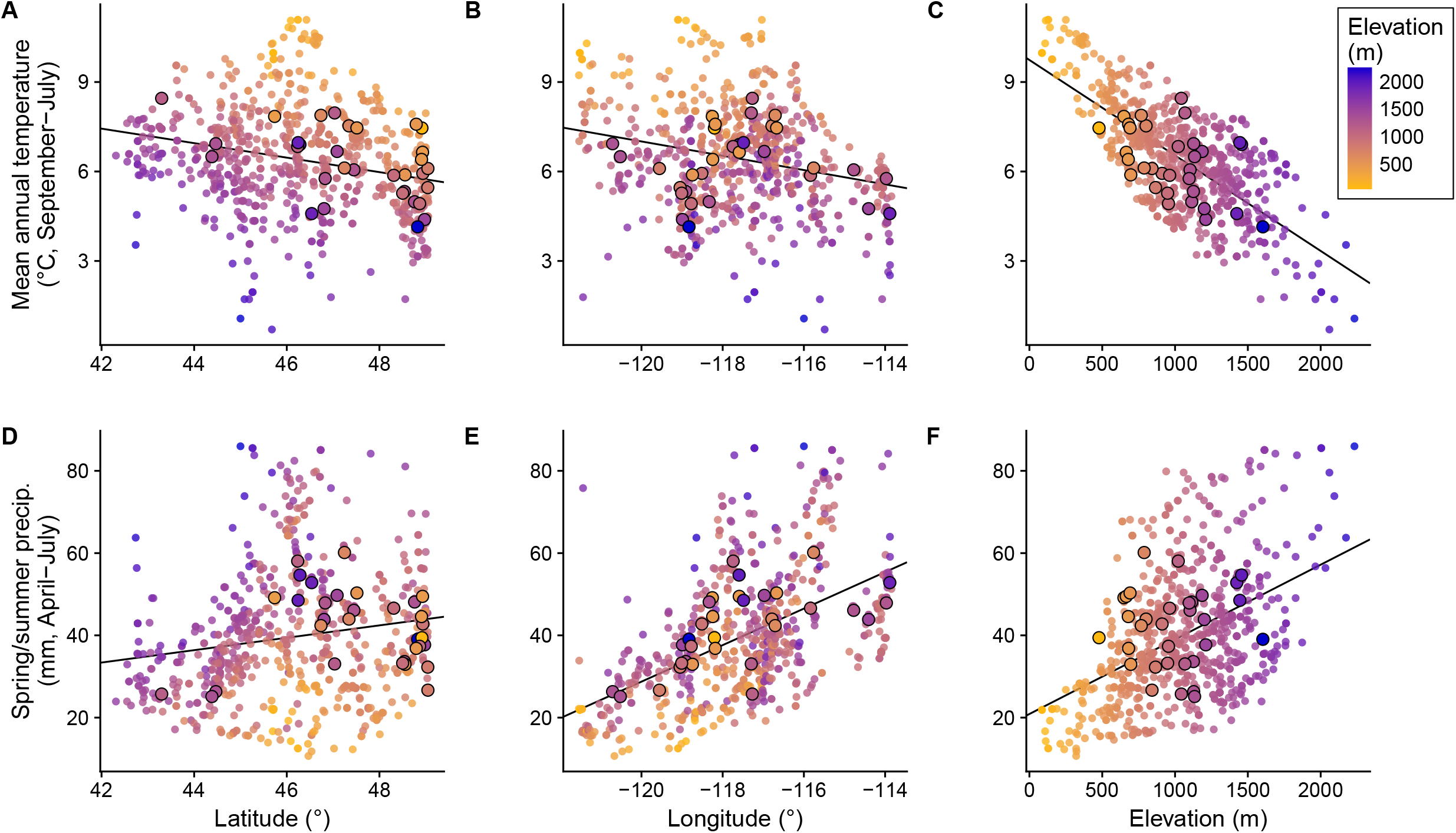
Relationships of climate and geography across the range of *Clarkia pulchella*. Small points represent all herbarium localities of *C. pulchella*, larger outlined points represent populations included in this study. Points are colored according to elevation. Temperature is influenced by **(A)** latitude, **(B)** longitude, and **(C)** elevation. Precipitation is also influenced by **(D)** latitude, **(E)** longitude, and **(F)** elevation. However, the interaction of these drivers results in climate that is heterogeneous across space. Climate data are 1951-1980 averages from PRISM (PRISM Climate Group, 2017). Trend lines are slopes from linear regression.

### Population selection, climate characterization, and seed collection

For this study, we selected populations that would allow us to decouple climatic and spatial axes of differentiation. For example, we wanted to include populations that were spatially near each other but climatically different and populations that were geographically distant but climatically similar. Monthly temperature and precipitation data from 1951-1980 for all populations were obtained from PRISM (PRISM Climate Group, 2017). We calculated the average temperature across the months that encompass the *C. pulchella* life cycle (September-July) and average precipitation when *C. pulchella* is most likely to be water-limited (April-July) for each population. Based on field observations and common garden trials (Bontrager and Angert, in prep), we considered these to be good candidates for variables that might have the potential to generate patterns of isolation by environment via selection against migrants. We first considered a set of 40 populations that we had located, then narrowed that set down to 32 populations that maximized variation in the relationship between spatial proximity and climatic similarity (Figure 1, Table S1). In July of 2014, we collected seeds from 12 plants separated by at least 0.5 m in each of those populations. Seeds from 17 populations were grown in the greenhouse beginning in December of 2014, and seeds from the remaining 15 populations were grown in growth chambers beginning in February of 2016.

### DNA Extraction

Tissue was harvested from the first cohort of plants in May 2015. Leaf or bud tissue was collected into 2 mL tubes on ice, then frozen at −80°C until DNA extraction. Tissue from the second cohort was collected onto dry ice in April 2016 and stored at −80°C until DNA extraction. DNA was extracted using DNeasy Plant Mini kits and DNeasy Plant 96 kits (Qiagen), following the protocol for frozen tissues. DNA extractions that did not have satisfactory 260/230 or 260/280 ratios were cleaned with ethanol precipitation. DNA was eluted and stored in 10mM Tris-HCl pH 8.

### Library preparation and sequencing

We prepared for two lanes of sequencing, with six individually barcoded samples from each population in each lane (191 or 192 individuals per lane, because we only had DNA of a high enough quality from a total 11 individuals from one population). Our library preparation protocol was a modified version of Poland et al. (2012). Libraries were prepared using 100 ng starting material per sample. DNA was digested in a 20 μL reaction using 8 units each of the enzymes MspI and Pst I-HF (New England Biolabs) in the supplied buffer. Digestion was carried out for 5 hours at 37°C, followed by 20 minutes at 65°C. Reactions were then stored overnight at 4°C. Ligation was performed in a 40 μL reaction in the same buffer as the digestion with 200 units of T4 DNA ligase (New England Biolabs) using 192 barcoded adapters and 12 common adapters on the opposite end. Ligation was performed for 3 hours at 22°C followed by a 20 minute hold at 65°C. Reactions were then cleaned with 1.6 volumes of SPRI beads and two 80% ethanol washes and resuspended in 12 μL of Tris-HCl pH 8.

Amplification was carried out in 10 μL reactions using 4 μL of cleaned ligation product, Kapa HIFI HotStart master mix (Kapa Biosystems), and primers from Poland et al. (2012). Amplification began at 98°C (30 s), followed by 14 cycles of 98°C (30 s), 62°C (20 s), 72°C (30 s), and a 72°C hold for 5 minutes. After amplification, samples were quantified using fluorometry, then each plate was pooled according to individual concentrations to yield a final product with equal amounts of library from each individual. This pooled library was run out on a 1.5% agarose gel and bands containing fragments 400 to 600 bp long were excised and cleaned using a gel extraction kit (Qiagen). The eluted product was cleaned and concentrated using SPRI beads.

Finally, we reduced the number of high copy fragments from our library using a protocol modified by M. Todesco from Shagina et al. (2010) and Matvienko et al. (2013). We began with 480 ng of each library in a 3 μL volume. To this we added 1 μL of hybridization buffer (200 mM HEPES pH 7.5, 2M NaCl, 0.8 mM EDTA), covered the reaction with mineral oil, heated it to 98°C for 2 minutes, then held it at 78°C for 3 hours. We then added 5 μL of duplex specific nuclease buffer (0.1 M Tris pH 8, 10mM MgCl_2_, 2mM DTT) and incubated at 70°C for 5 minutes. We then added 0.2 μL of duplex specific nuclease and incubated at 70°C for another 15 minutes, then stopped the reaction with 10 μL of 10 mM EDTA. We then reamplified the library using the same reagents as above in a 25 μL reaction with 2-4 μL of template and cleaned again with SPRI beads. Libraries were stored at −20°C until sequencing. Libraries were sequenced with paired-end 100 bp reads on the Illumina HiSeq 2000 platform at the Biodiversity Research Centre at UBC.

### Alignment and SNP calling

Sequences were processed and aligned using components of the Stacks pipeline (version 1.40, Catchen et al., 2011, 2013). Reads with uncalled bases or low quality scores (average quality in a 14-base sliding window *<*10) were discarded. Ten samples had far fewer reads than the rest and these were excluded prior to alignment. Paired end reads were pooled with first end reads, i.e. during alignment and SNP detection the two ends of each read were treated as if they were independent loci (we later checked for linkage disequilibrium among SNPs). During initial “stacking” and catalog building we allowed sequences to diverge at 3 bases, and set the minimum depth of coverage required to create a stack at 3 (Rochette and Catchen, 2017). Modifications to these parameters did not result in substantial differences in values of pairwise F_ST_ (data not shown). The maximum number of stacks per locus was set to 3, and gapped alignments were not allowed. We enabled the removal algorithm, which drops highly repetitive stacks (removes initial stacks that have *>*2 SD coverage relative to individual sample mean), and the deleveraging algorithm, which breaks up or removes over-merged sequences. Our catalog was built using all samples. We employed the rxstacks corrections module to correct or omit loci with putative sequencing errors, loci with low log-likelihoods (*<*-10), confounded loci, and loci with excess haplotypes.

SNP tables were generated using the populations module of Stacks. Initial inspection of PCA plots using SNPRelate (Zheng et al., 2012) revealed three individuals that were not clustering with the other individuals from their populations. We consider it more plausible that these represent mis-labeled samples in the field, greenhouse, or lab than long-distance migration events. Downstream analyses were performed without these individuals. Therefore, in our final dataset, seven populations had only 11 individuals, one population had only 10, one population had only 8, and the remaining 23 populations were each represented by 12 individuals. In our analyses we included only loci that had coverage of at least 12x in 75% of individuals in 75% of populations, with a minimum minor allele frequency of 0.05 and a maximum heterozygosity of 70% across all populations. We checked that pairwise F_ST_ was not sensitive to these parameter choices. In case of multiple SNPs occurring in a single locus, we kept just the first one. After applying these filters, 2983 SNPs were retained. Linkage disequilibrium was generally low among our loci (r^2^ *<*0.2 for 99.9% of pairs of SNPs). F_ST_ was calculated using the implementation of Weir and Cockerham (1984) and expected heterozygosity (within-population gene diversity) was calculated using methods from Nei (1987) in the R package hierfstat (Goudet and Jombart, 2015). Because populations varied in the average proportion of loci that were successfully genotyped (three populations had *<*60% success; among all populations the median success rate was 78% and the range was 23-92%), we checked that expected heterozygosity did not correlate with genotyping success rate (r = 0.27, *P* = 0.13).

### Quantifying isolation by environment vs. isolation by distance

We used BEDASSLE (Bradburd et al., 2013) to estimate the relative contributions of geographic distance and climatic differences to genetic differentiation. BEDASSLE is implemented in R (R Core Team, 2017), and it employs a Markov chain Monte Carlo (MCMC) algorithm to estimate the relative effect sizes of geographic distance and environmental differences on covariance in allele frequencies among populations. As environmental covariates, we used pairwise differences in average September-July temperature and average spring/summer precipitation (April-July). We initially generated resistance-weighted distances between populations using projected habitat suitability as a conductance matrix, but these distances were highly correlated with actual geographic distances and did not produce better model fits in preliminary analyses, so we did not use them in these models. We estimated effect sizes of geography, temperature, and precipitation differences using all 32 populations, but also ran BEDASSLE for subsets consisting of populations clustered in the central and northern parts of the range (indicated in Table S1) to see if we could detect effects of the environment that may be obscured or weakened at large geographic scales. Prior to analysis, we divided pairwise geographic distance and precipitation differences by their standard deviations so that these predictors were on a scale more similar to pairwise temperature differences. We ran these models for 10 million generations, and thinned the chains by sampling every 1000 generations. We visually inspected MCMC traces and marginal distributions to ensure that models reached stationary distributions. All results are reported after a burn-in of 20%, with effect sizes back-transformed to the scale of the original data. We checked these results against partial Mantel tests of pairwise geographic, temperature, and precipitation differences on pairwise F_ST_ using the R package phytools (Revell, 2012). We did not rely upon partial Mantel tests as our main analytical method because of their potential to have inflated Type I error rates (Guillot and Rousset, 2013).

### Assessment of spatially continuous vs. discrete genetic differentiation

We were interested in evaluating whether population structure was well-described by modelling populations as admixtures between multiple discrete genetic groups, as might be caused by geographic barriers (i.e., the Rocky Mountains) or historic phylogeographic processes. We evaluated how well models prescribing various numbers of discrete genetic groups described differentiation and similarity among our populations using conStruct (Bradburd et al., 2017). conStruct is implemented in R (R Core Team, 2017), and is similar to the frequently-used program Structure (Pritchard et al., 2000) but allows genetic differentiation to increase with geographic distance between populations even when these populations draw from the same genetic groups. In the spatial implementation of this program, populations are composed of admixture from a user-specified number of discrete layers (K), and genetic similarity decays with geographic distance within each of these layers. We ran conStruct for 1000 iterations setting the number of layers to 1, 2, 3, 4, and 5. We compared the fits of each of these different parameterizations using cross-validation and by evaluating the contribution of each additional layer to the total covariance of these loci. For cross-validation, we fit models with subsets containing 90% of loci and evaluated the resulting model fit by calculating the log likelihood of the remaining loci. We performed 100 replicate cross-validation runs.

### Exploring spatial patterns in genetic diversity

We examined whether population genetic diversity (as estimated by expected heterozygosity) exhibited geographic trends. We used linear models in R (R Core Team, 2017) to test whether expected heterozygosity was predicted by latitude or by proximity to the range edge (as measured by the distance of a population to the nearest edge of a polygon drawn around all localities of the species; Figure 1).

## Results

### Isolation by environment vs. geographic distance

Overall F_ST_ among these populations is 0.135. Genetic differentiation between populations of *Clarkia pulchella* is primarily structured by geographic distance, with no apparent contribution of the environmental variables that we have considered here (Figure 3). The effect size of a temperature difference of one degree (C) relative to the effect of 100 km of geographic distance is 1.18 × 10^−7^ (95% credible interval = 8.52 × 10^−8^ - 1.58 × 10^−7^; Figure S1A), and the effect of 10 mm of spring/summer precipitation difference relative to the effect of 100 km of geographic distance is 5.84 × 10^−7^ (95% credible interval = 1.50 × 10^−8^ - 2.98 × 10^−6^; Figure S1B). The scales at which these ratios are presented are arbitrary, but they were chosen so that the range of values among populations is on the same order of magnitude: 100 km represents about one sixth of the maximum pairwise geographic distance, 1°C represents approximately one fourth of the maximum pair-wise temperature difference, and 10 mm precipitation represents about one fourth of the maximum pairwise precipitation difference. The climatic effect sizes we found are so small that the effects of these variables can be considered nonexistent in terms of their biological importance. Effects of environmental differences did not emerge at smaller geographic scales in subsets of populations in the north (effect of temperature differences relative to geographic distance: 5.89 × 10^−8^ (8.61 × 10^−9^ - 1.14 × 10^−7^), effect of precipitation differences relative to geographic distance: 9.73 × 10^−6^ (5.81 × 10^−7^ - 2.11 × 10^−5^); Figure S2) or center (effect of temperature: 2.34 × 10^−7^ (1.44 × 10^−8^ - 4.73 × 10^−7^), effect of precipitation: 9.46 × 10^−7^ (3.06 × 10^−8^ - 4.80 × 10^−6^); Figure S3). These conclusions are consistent with the results of partial Mantel tests, in which only pairwise geographic distance is a significant predictor of pairwise F_ST_ (Table 1).

**Figure 3.**
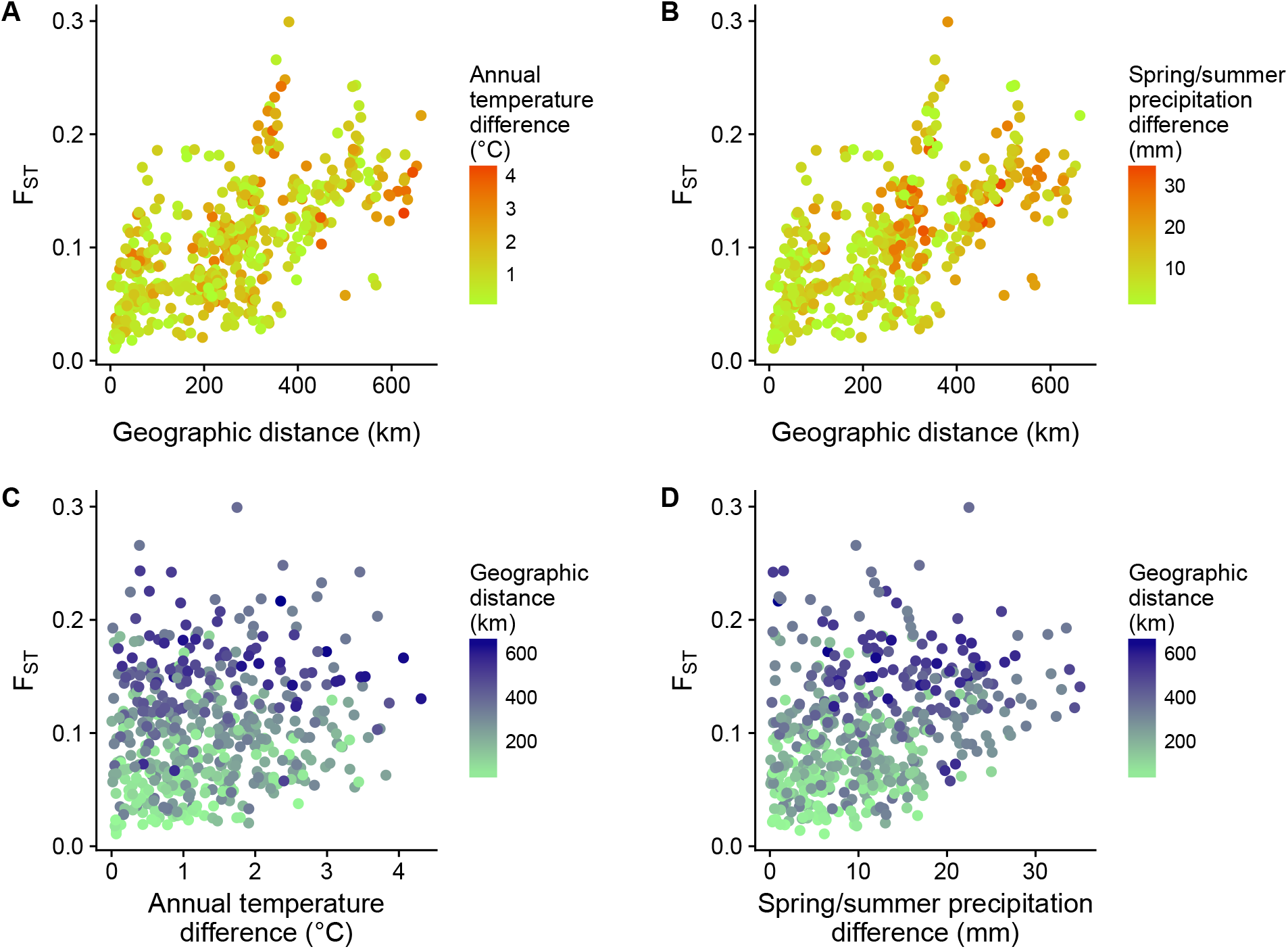
Pairwise genetic differentiation (F_ST_) of populations of *Clarkia pulchella* increases with geographic distance (x-axis in **A** and **B**), but shows no discernible relationship to temperature differences (color in **A**) or precipitation differences (color in **B**). An alternative visualization is presented in **(C)** and **(D)**, in which climate differences are plotted on the x-axis and geographic distance is indicated with color. Climate data are 1951-1980 averages from PRISM (PRISM Climate Group, 2017).

**Table 1.** Results of partial Mantel tests of pairwise geographic distance (km), pairwise temperature differences (°C, September-July, 1951-1980 averages), and pairwise precipitation differences (mm, April-July, 1951-1980 averages) on pairwise genetic differentiation (F_ST_) among populations of *Clarkia pulchella*. Climate data are 1951-1980 averages from PRISM (PRISM Climate Group, 2017).

| Region | $R^2$ | $P$-value | Predictor | Coefficient | t-statistic | $P$-value |
| --- | --- | --- | --- | --- | --- | --- |
| Entire range | 0.42 | **0.001** | Geographic distance | 0.0002 | 15.73 | **0.001** |
|  |  |  | Temperature differences | 0.0028 | 1.44 | 0.486 |
|  |  |  | Precipitation differences | 0.0006 | 2.46 | 0.209 |
| North | 0.36 | **0.008** | Geographic distance | 0.0006 | 4.65 | **0.006** |
|  |  |  | Temperature differences | 0.0061 | 1.63 | 0.377 |
|  |  |  | Precipitation differences | 0.0004 | 0.57 | 0.692 |
| Center | 0.44 | 0.06 | Geographic distance | 0.0006 | 4.87 | **0.001** |
|  |  |  | Temperature differences | 0.0111 | 0.56 | 0.463 |
|  |  |  | Precipitation differences | -0.0005 | -0.45 | 0.737 |

### Genetic structure of populations

The genetic structure of these populations is explained slightly better by a model of admixture between two genetic groups than by a model of continuous genetic differentiation across space, as indicated by the increase in predictive accuracy in models where two layers were allowed rather than one (Figure 4). Northern populations primarily belong to one genetic group, while southern populations belong to another, and populations from mid-latitudes are a mix of the two (Figure 5). Allowing more than two layers did not improve predictive accuracy (Figure 4). Note that populations east of the Rocky Mountains (populations D9, D10, and P12) never formed a separate group, regardless of the number of layers allowed (results not shown). Although models with two layers did have greater predictive accuracy than those with one, when K = 2 the amount of covariance contributed by the second layer was small relative to the first (Table 2).

**Figure 4.**
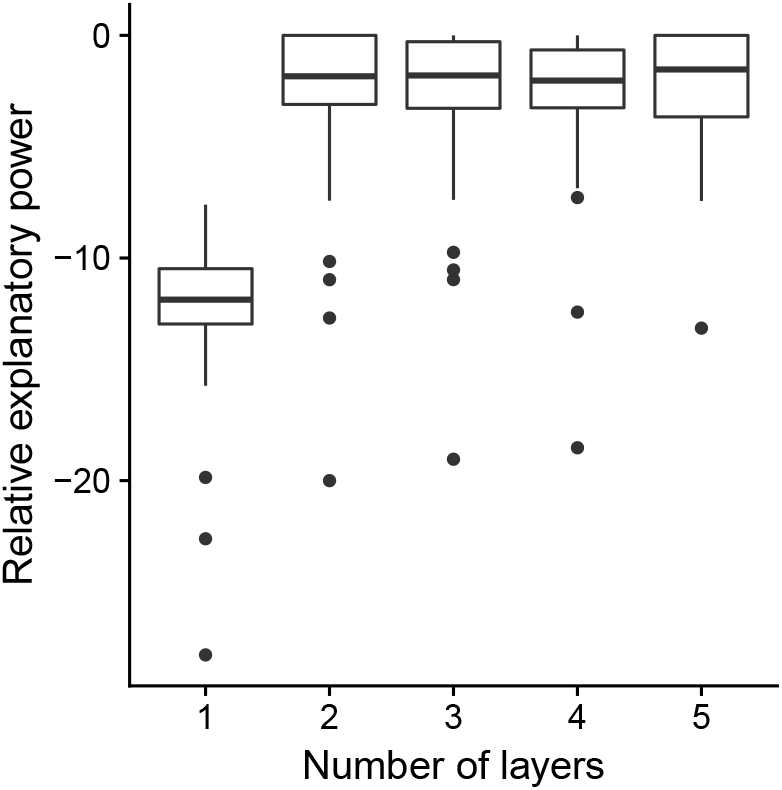
Results of 100 replicate cross-validation runs of conStruct with the number of layers set to 1, 2, 3, 4, or 5. In each replicate, the model is built using 90% of loci, and the log-likelihood of the remaining loci is calculated. Predictive accuracy is then calculated as the difference in log-likelihood between each model and the best model (i.e. the best number of layers) in each replicate. These results indicate that models constructed with two layers are best, because they provide as much explanatory power as other models without further complexity.

**Figure 5.**
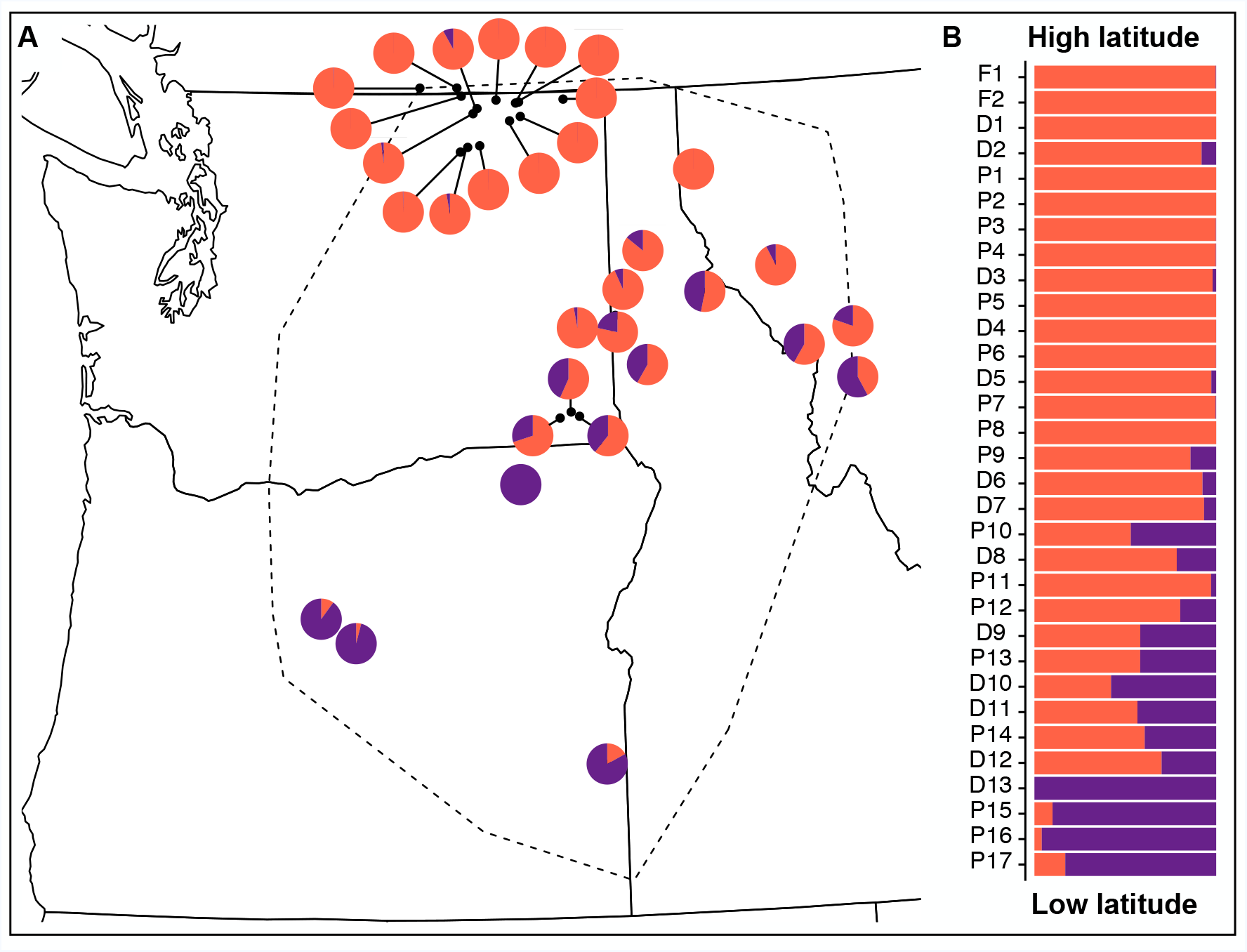
Admixture proportions of each of 32 populations of *Clarkia pulchella* estimated from by conStruct with K = 2. **A** Admixture proportions are shown in geographic space and **(B)** arranged by latitude. Population ID codes are consistent with Table S1 and Figure 1.

**Table 2.** Covariance contributions of each layer in conStruct models with the number of layers (K) set to 1, 2, 3, 4, or 5.

| Number of layers | 1 | 2 | 3 | 4 | 5 |
| --- | --- | --- | --- | --- | --- |
| Layer contributions | 1.000 | 0.9004 | 0.8062 | 0.8014 | 0.8925 |
|  | - | 0.0996 | 0.1043 | 0.1541 | 0.0795 |
|  | - | - | 0.0895 | 0.0438 | 0.0204 |
|  | - | - | - | 0.0007 | 0.0055 |
|  | - | - | - | - | 0.0021 |

### Geographic trends in genetic diversity

Genetic diversity increases with latitude among these populations (estimate = 0.0104, SE = 0.0019, df = 30, *P <* 0.0001, Figure 6A), but is not related to distance from the range edge (df = 30, *P* = 0.811). Genetic diversity appears to be lower in populations in the southern half of the range, and also in populations near the eastern range edge, but is higher in central and northern populations (Figure 6B).

**Figure 6.**
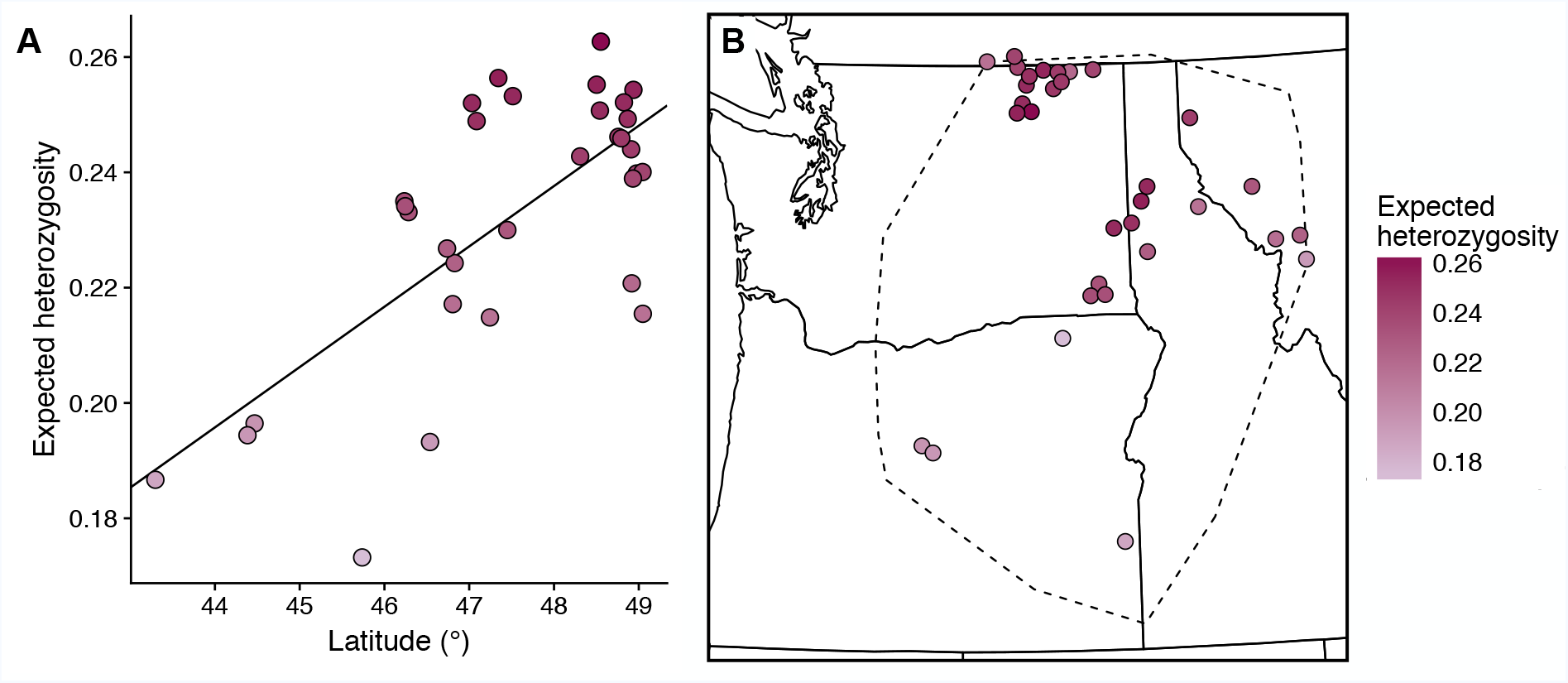
**(A)** Expected heterozygosity increases with latitude across the range of *Clarkia pulchella*. **(B)** Expected heterozygosity appears to be higher in central and northern parts of the range, but lower in the south and east.

## Discussion

We contrasted the relative effects of geographic vs. climatic distances on genetic differentiation in *Clarkia pulchella*, examined whether geographic structure in this species could be described by assigning populations to distinct genetic groups, and tested for geographic gradients in genetic diversity. Our analyses revealed a genetic structure that is predominantly shaped by geographic distances between populations. In addition to this pattern of isolation by distance, populations partition into northern and southern groups, with admixed populations in the center of the range. Genetic diversity was highest in northern and central populations, resulting in a trend of increasing genetic diversity with latitude.

### Populations of *Clarkia pulchella* are isolated by distance

At the scale of the geographic range in *Clarkia pulchella*, isolation by distance is the dominant pattern. This likely reflects gene flow that is strongly restricted by geographic distances between populations. This is perhaps not surprising, given that this species has no obvious mechanism for seed dispersal and our best guess is that gene flow between populations is facilitated by occasional pollen movement by bumblebees, hummingbirds, and other floral visitors. In the case of an absence of climatically structured seed and pollen movement, selection against migrants and their offspring is the remaining mechanism that could drive isolation by environment. While *C. pulchella* does appear to be locally adapted to historic climate (Bontrager and Angert, in prep), selection against foreign genotypes may not be strong enough to preempt the spread of neutral loci, even as recently-arrived loci that confer poor performance in a given environment are purged. This could lead to a signal of isolation by distance at neutral loci, while populations are still adaptively differentiated based on their local climate.

It is possible that the absence of an effect of temperature and precipitation differences on genetic structure is the result of our experimental design, and that environmental differences might matter in other contexts. There may be environmental variables other than those we have considered here that are more important in determining the movement of genes or the realized rate of gene flow among populations. These could be climatic, but also could include soil characteristics, or local adaptation to competitors, pollinators, or soil biota. It is also possible that the effects of environmental differences are more detectable at smaller spatial scales. For example, in some plant species, differences in phenological timing along local snowmelt gradients structure gene flow to a greater extent than geographic distances (Hirao and Kudo, 2004; Shimono et al., 2009). Similar processes may play out in *C. pulchella* as well, possibly along local elevation gradients.

### Populations are admixtures of northern and southern genetic groups

Rather than mountain ranges separating populations into genetic groups, we detected underlying population structure that divides the species into northern and southern groups, with admixed populations in the middle. This suggests that perhaps the Columbia Basin, a low-elevation, relatively flat area in south-central Washington (Figure 1), is a barrier to gene flow in this species. Species distribution models indicate that it is an area of low suitability (Bontrager and Angert, 2016) and few occurrences of *Clarkia pulchella* have been recorded in this region. Most studies of population genetic structure in the Pacific Northwest focus on mesic forest species that occupy the wet western slopes of both the coastal and Rocky Mountains (Shafer et al., 2010), and these studies often find differentiation between western and eastern populations. Phylogeographic research on species occupying the arid inter-mountain region is less common. In the Great Basin pocketmouse, a species with a range that overlaps with that of *C. pulchella*, a north-south split in genetic structure was detected in approximately the same location as in our results (Riddle et al., 2014). It is possible that the Columbia Basin (or some geographic feature within it) represents a barrier to gene flow, either past or ongoing, for a variety of taxa that occupy the dry intermountain region. The habitat affinity of species can influence the effect of glaciation events on genetic structure (Massatti and Knowles, 2014), therefore further work on *C. pulchella*, including paleoclimate modelling or modelling demographic history, might allow for an interesting contrast with the relatively well-studied mesic flora of the Pacific Northwest.

### Genetic diversity increases with latitude

We expected we would see lower genetic diversity at higher latitudes, but we detected the opposite: genetic diversity was highest in north-central and northern populations (though the total magnitude of variation in expected heterozygosity was not large). This latitudinal pattern is somewhat surprising, because northern populations are in areas that were under glaciers during the last glacial maximum, and we expected that range expansion into this area after their retreat would result in a signature of lower genetic diversity. When high levels of genetic diversity are present in areas of past range expansion, this can sometimes be attributed to the mixing of populations that had previously been persisting in multiple refugia (Petit et al., 2003; Brunsfeld and Sullivan, 2005). Species in the northern Rocky Mountains that are presumed to have occupied multiple refugia often exhibit some degree of contemporary differentiation between northern and southern populations (Brunsfeld et al., 2001; Brunsfeld and Sullivan, 2005), a pattern consistent with what we have found in *Clarkia pulchella*. Regardless of the location or number of refugia that *C. pulchella* previously occupied, it is also possible that range expansion was not accompanied by reductions in genetic diversity in this species, as is sometimes the case in other systems (Vandepitte et al., 2017).

The more common expectation for geographic patterns in genetic diversity is that range edge populations will have lower genetic diversity (Vucetich and Waite, 2003). This prediction is based on the assumption of an abundant center distribution pattern, in which edge populations are small, and may experience frequent turnover or constant directional selection (if they are far from the phenotypic optima of an extreme environment). Our results are not consistent with this being the case for *C. pulchella*, at least not at all range edges. We note however that populations at southern and eastern edges do appear to have lower genetic diversity relative to the northern and north-central populations, and further work could be done to investigate the processes that might generate this pattern.

### Conclusions

Our investigation of the genetic structure of *Clarkia pulchella* has revealed some intuitive patterns, as well as surprising ones. Despite substantial heterogeneity in climate across the species’ range, genetic similarity is primarily determined by geographic proximity. Though a signal of isolation by distance is not surprising in a sessile organism studied at a large spatial scale, the absence of any effect of environment indicates that to the extent that populations experience gene flow, it may be from both similar and divergent environments. This species does not exhibit geographic patterns of genetic diversity consistent with our expectations for a recently expanded northern range edge nor a range limited by adaptation. These results would be complemented by future work examining mechanisms of contemporary gene flow and historic demographic processes in *Clarkia pulchella*.

## Acknowledgements

We would like to thank C. Caseys and M. Todesco for their generous guidance and training during library preparation. Library preparation protocols were optimized by M. Todesco, K. Ostevik, and B. Moyers in the Rieseberg Lab at the University of British Columbia. G. Owens provided helpful advice on bioinformatics methods. E. Fitz assisted with locating populations of *C. pulchella* in the field and A. Wilkinson assisted with plant cultivation. We appreciate thoughtful comments on a draft of this manuscript from S. Aitken, M. Whitlock, and J. Whitton. Funding for this project was provided by the Washington Native Plant Society, the Botanical Society of America and the Botanical Society of America Genetics Section, and a National Sciences and Engineering Research Council Discovery Grant to ALA. MB was supported by a University of British Columbia Four-year Fellowship.

## Author contributions

MB conceived of the project in consultation with ALA. MB performed all field work, lab work, bioinformatics, and analyses with guidance from ALA. MB generated all figures and tables and wrote the manuscript with frequent conversation and comments from ALA.

## Data accessibility

Data and code are hosted on Github at https://github.com/meganbontrager/clarkia-pulchella-popgen and will be archived on Dryad or a similar repository upon publication. Sequences are archived on the NCBI Sequence Read Archive (SUB4307183).

---

**Table S1.**
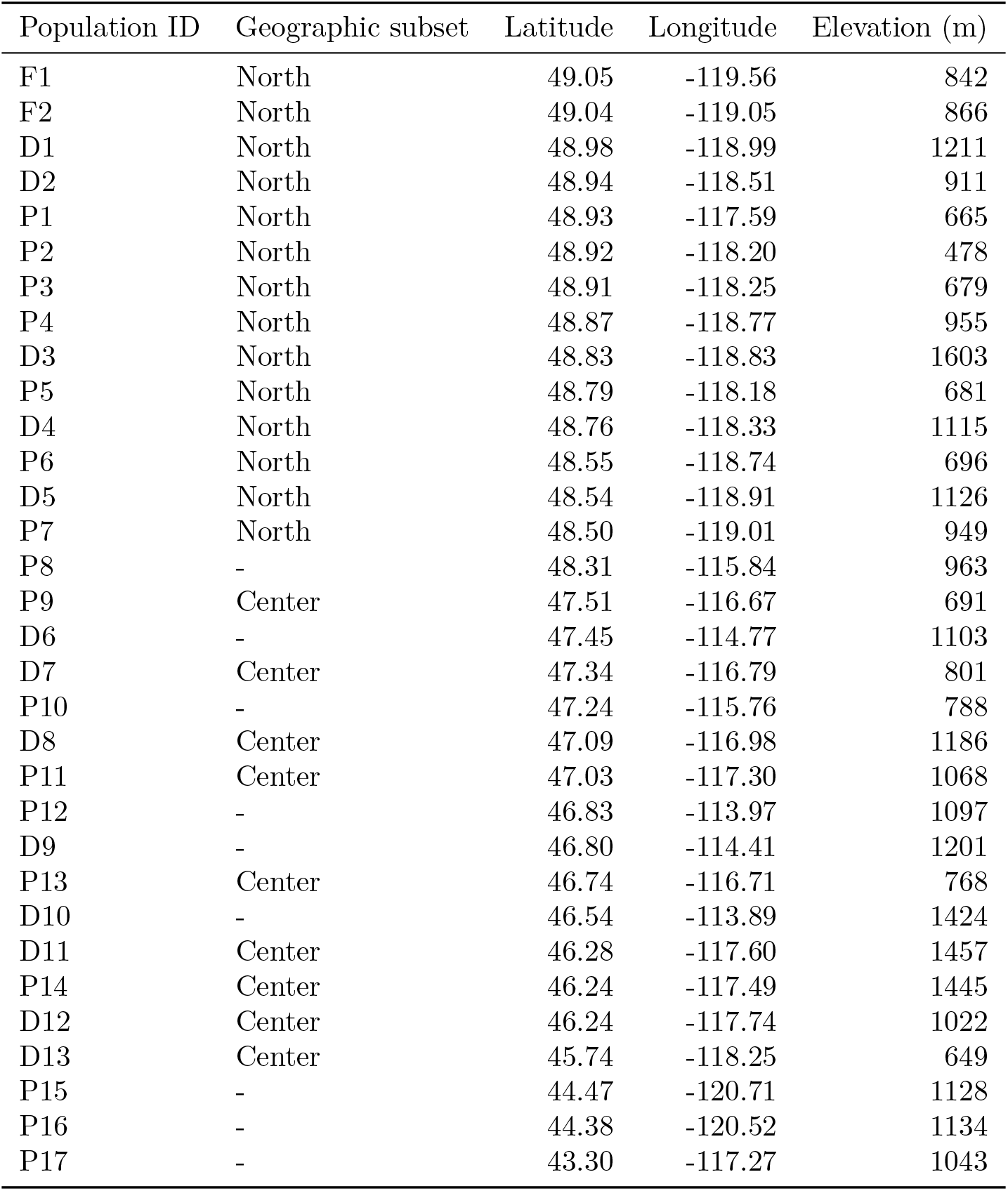
Geographic locations and elevations of populations of *Clarkia pulchella* included in these analyses. Population IDs are consistent with Figure 1. The populations included in analyses of geographic subsets are indicated.

**Figure S1.**
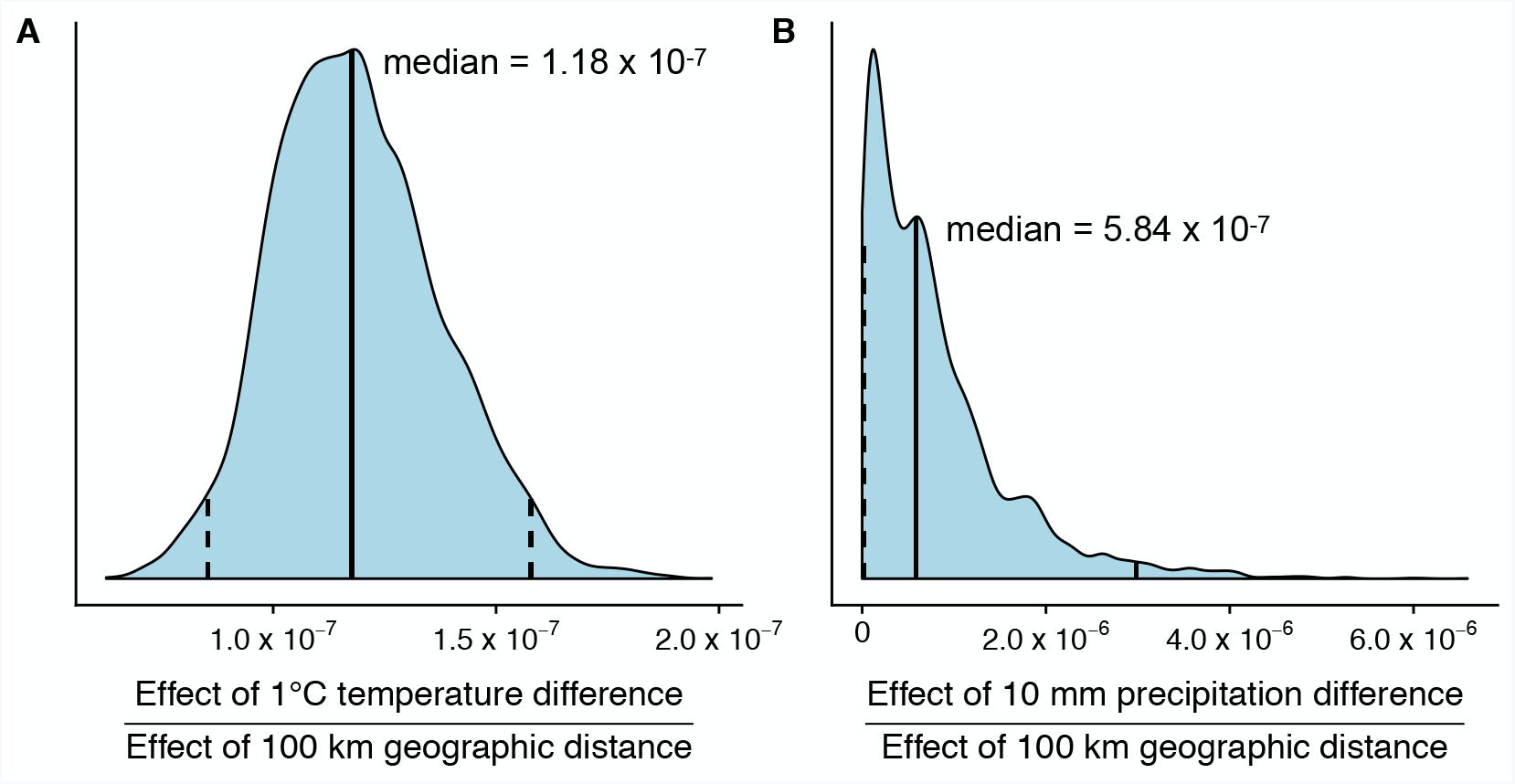
Marginal posterior distributions, median values (solid lines) and 95% credible intervals (dashed lines) of the ratio of the effect sizes of **(A)** temperature vs. geographic distance and **(B)** spring/summer precipitation vs. geographic distance on genetic differentiation of populations of *Clarkia pulchella* after a burn-in of 20%.

**Figure S2.**
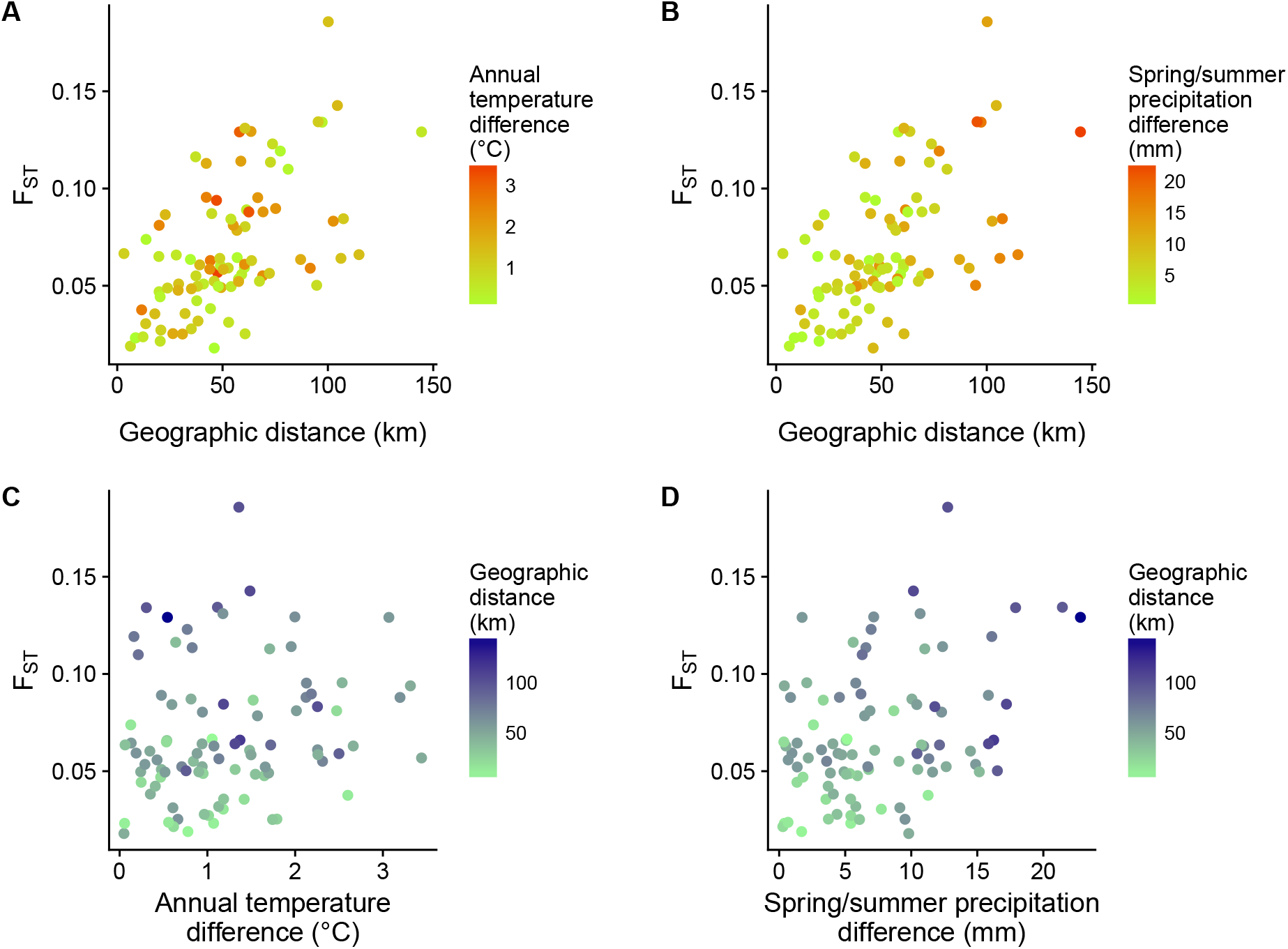
Relationship between pairwise geographic distance (x-axis in **A** and **B**), temperature differences (color in **A**) or precipitation differences (color in **B**), and genetic differentiation (F_ST_) among populations in the northern part of the geographic range of *Clarkia pulchella*. An alternative visualization is presented in **(C)** and **(D)**, in which climate differences are plotted on the x-axis and geographic distance is indicated with color. Climate data are 1951-1980 averages from PRISM (PRISM Climate Group, 2017).

**Figure S3.**
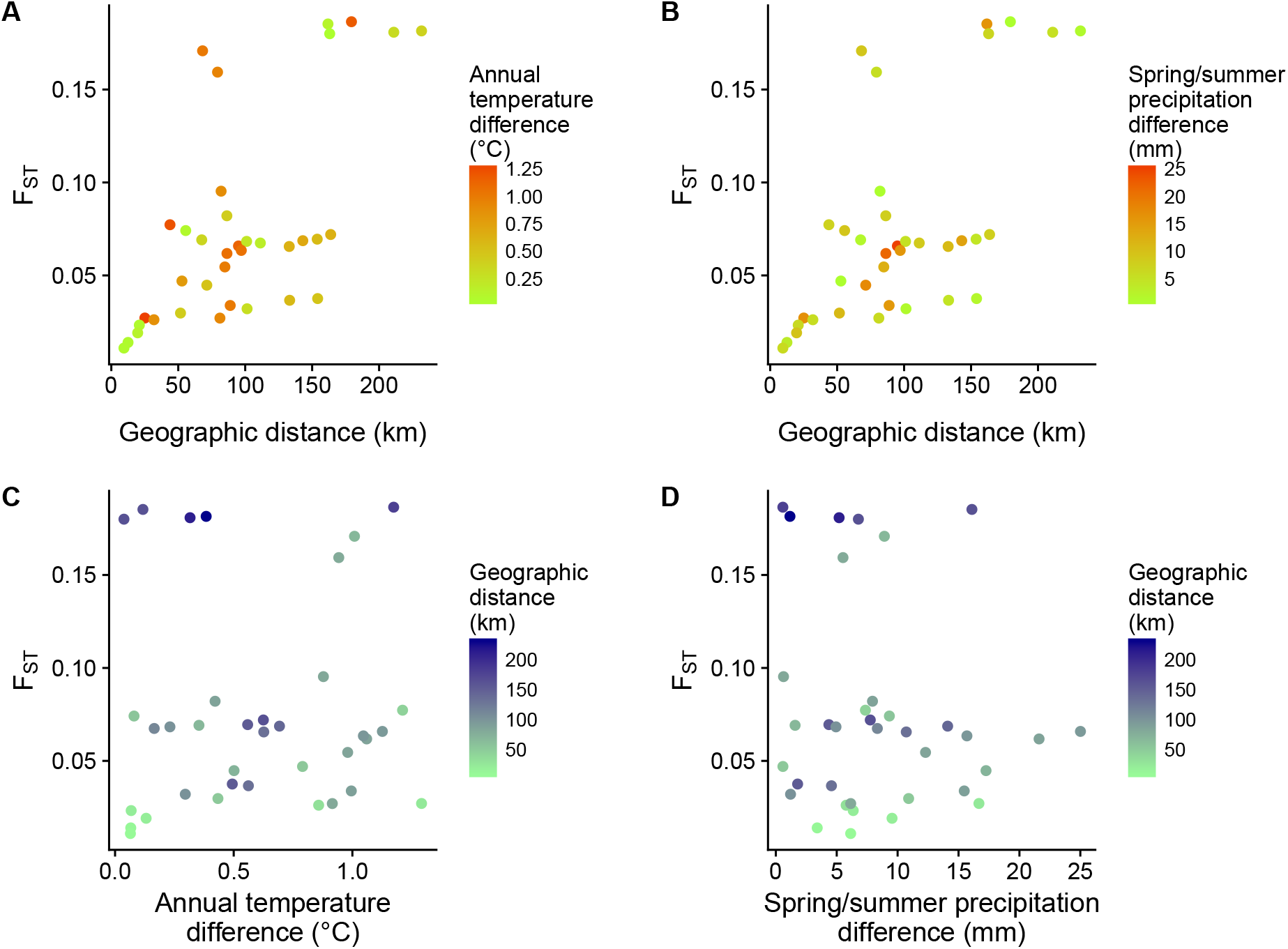
Relationship between pairwise geographic distance (x-axis in **A** and **B**), temperature differences (color in **A**) or precipitation differences (color in **B**), and genetic differentiation (F_ST_) among populations in the central part of the geographic range of *Clarkia pulchella*. An alternative visualization is presented in **(C)** and **(D)**, in which climate differences are plotted on the x-axis and geographic distance is indicated with color. Climate data are 1951-1980 averages from PRISM (PRISM Climate Group, 2017).

